# Conversion of aromatic compounds from fractionated industrial hydrolysis lignin by *Pseudomonas putida* and environmental microbial strains

**DOI:** 10.1101/2023.10.19.563119

**Authors:** Philip A. Morehead, Henry Vider, Christina Mürk, Signe Viggor, Merike Jõesaar, Scott Bottoms, Siim Salmar, Maia Kivisaar, Mart Loog

**Author notes:** Corresponding authors. E-mail address (S. Bottoms); (S. Salmar).

## Abstract

**Background:** The utilization of *Pseudomonas putida* was explored in this study as a promising approach for lignin valorization. To this end, dry hydrolysis lignin was used as a feedstock for the first time. Hydrolysis lignin is a product of the enzymatic hydrolysis and separation of cellulose and hemicellulose from the lignin backbone in diverse lignocellulosic sources. Various fractionation techniques were applied to obtain lignin monomers and multimers in solution for use as a growth medium for *P. putida*, whose tolerance of inhibitory phenolic compounds distinguishes it from most bacteria.

**Results:** Physiological evaluations revealed that *Pseudomonas putida* strains KT2440 and PaW85 exhibited broad pH tolerance ranges, with robust growth observed at elevated pH levels. Batch fermentations using hydrolysis lignin (HL) solutions showed complete consumption of sugars within 24 hours, demonstrating the viability of fractionated HL as a substrate for *P. putida* cultivation. HPLC analysis of HL monomer concentrations during simulated fed-batch fermentation revealed rapid catabolism of catechol and increased CCMA concentration, followed by stabilization, indicating that CCMA is synthesized more quickly than degraded when the initial catechol concentration is high. Filtered alkaline HL fractionations yielded more than twice as much catechol as unfiltered fractionations. Screening of indigenous bacterial strains isolated from various soil and water samples (CELMS Collection, website http://eemb.ut.ee) identified five new candidate strains for CCMA production, two for PCA production, and three for vanillic acid production.

**Conclusions:** The novel use of fractionated hydrolysis lignin as a growth medium shows potential for lignin valorization and chemical production. Filtered alkaline fractionation yields more catechol and is superior for *cis,cis*-muconic acid production; however, unfiltered fractionations may be more suitable for other compounds and upscaling. Further investigation of screened strains could reveal more efficient enzymes, which could be optimized and transformed into *P. putida* in future research.

## Introduction

The most critical global environmental issue is the rising concentration of atmospheric greenhouse gases (GHG), which leads to climate change due to increased global temperatures [1]. Since *circa* 1750, the atmospheric CO_2_ concentration has steadily risen from around 277 ppm to approximately 410 ppm in 2019 [2]. As the impact of climate change continues to raise environmental concerns, it is crucial to address plastic emissions. The environmental impacts of plastics production, particularly GHG emissions, have received minor scrutiny [3]. However, plastics emissions have doubled between 1995 and 2015, amounting to 4.5% of global carbon emissions [4].

Lignin shows growing potential to mitigate the climate impact of petrochemical production as it is the only abundant renewable feedstock comprised of polymerized aromatic compounds. Lignocellulosic biomass is a widely used resource on a global scale, contributing to the production of bioenergy and various value-added products [5]. Traditionally, lignin was considered a low-value waste product; however, recent advancements have revealed the potential to convert it into value-added products [6]. Lignin valorization involves depolymerizing and upgrading lignin subunits into value-added products, which is crucial in achieving environmental sustainability and economic viability for various applications [7,8].

*Pseudomonas putida* is an excellent candidate for industrial biocatalysis, given its biochemical versatility, and has significantly contributed to various areas of research, including biotechnological applications [9]. *P. putida* is distinguished from other microorganisms by naturally possessing a remarkable array of biochemical functions and adaptable metabolisms [10]. These include catabolic pathways for various carbon, nitrogen, and phosphorus sources and rich secondary metabolisms intricately connected to a robust core biochemistry [9].

The bacterial production of *cis,cis*-muconic acid (CCMA) using lignin-based carbon sources is gaining attention as CCMA can be used as a precursor for the synthesis of adipic acid and terephthalic acid, which are raw materials for plastics production [11]. Through the benzoate degradation pathway (BDP), protocatechuic acid (PCA) and catechol can be produced from lignin-derived aromatic precursors. PCA and catechol can be further transformed into β-carboxymuconate and CCMA by protocatechuate 3,4-dioxygenase and catechol 1,2-dioxygenase [12]. Deploying molecular biology tools in *P. putida* along the BDP can potentially increase the production of target compounds.

## Methods

### Bacterial strains

Bacterial strains used in this study were obtained from the Collection of Environmental and Laboratory Microbial Strains (CELMS); financed by the Estonian Ministry of Education and Research (RLOMRCELMS), the public catalog of which is available on the Estonian Electronic Microbial Database (EEMB) website http://eemb.ut.ee (**Tab. 1)**. *Mitsuaria chitosanitabida* 1R2A4, *Pseudomonas putida* 1S7, *Pseudomonas fluorescens* PC20, *Pseudomonas thivervalensis* P101, *Pseudomonas syringae* KM2R4, *Pseudomonas stutzeri* VMR7, and *Arthrobacter sulfonivorans* Phe16 were kindly donated by Eeva Heinaru (Institute of Molecular and Cell Biology, Tartu, Estonia). *Enterobacter* sp./*Lelliottia* sp. T13, *Pseudomonas* sp./*defluvii, Pseudomonas putida* PJK7, *Pseudomonas stutzeri/xanthomarina* T10, *Serratia proteamaculans* SM, and *Rhodococcus* sp./*erythropolis* were kindly donated by Merike Jõesaar (Institute of Molecular and Cell Biology, Tartu, Estonia). *Pseudomonas resinovorans* A21, *Rhodococcus erythropolis* L19, *Rhodococcus erythropolis* A26, *Rhodococcus erythropolis* A15, *Rhodococcus erythropolis* A8, *Cellulomonas* sp. A11B, *Bacillus subtilis* A4X, *Bacillus subtilis* A24, *Bacillus simplex* L12ABA, and *Bacillus pumilus* A7 were kindly donated by Elerin Toomik (Tasmanian Institute of Agriculture, Hobart, Australia). *Kocuria rosea* 43SK1 was kindly donated by Eeva Heinaru, Sulev Kuuse, and Julius Tarand (Institute of Molecular and Cell Biology, Tartu, Estonia). *Acinetobacter johnsonii* 6S4 was kindly donated by Eeva Heinaru and Sulev Kuuse (Institute of Molecular and Cell Biology, Tartu, Estonia).

**Table 1.** Strains used in this study. The strains were obtained from the Collection of Environmental and Laboratory Microbial Strains (CELMS); financed by the Estonian Ministry of Education and Research (RLOMRCELMS), the public catalog of which is available on the Estonian Electronic Microbial Database (EEMB) website http://eemb.ut.ee

| Strains | Species | Relevant Genotype | Source/Reference/Isolated by |
| --- | --- | --- | --- |
| PaW85 | *Pseudomonas putida* | 17 kb transposon Tn4652 in chromosome. Plasmid-free strain | Bayley *et al.*, 1977 [13] |
| KT2440 | *Pseudomonas putida* | Plasmid-free derivative of *Pseudomonas putida* mt2 | Regenhardt *et al.*, 2002 [14] |
| 1R2A4 | *Mitsuaria chitosanitabida* |  | Eeva Heinaru (CELMS) |
| 1S7 | *Pseudomonas putida* |  | Eeva Heinaru (CELMS) |
| T13 | *Enterobacter* sp./*Lelliottia* sp. |  | Merike Jõessar (CELMS) |
| VAS4 | *Pseudomonas* sp./*defluvii* |  | Merike Jõessar (CELMS) |
| PJK7 | *Pseudomonas putida* |  | Merike Jõessar (CELMS) |
| T10 | *Pseudomonas stutzeri*/xanthomarina |  | Merike Jõessar (CELMS) |
| SM | *Serratia proteamaculans* |  | Merike Jõessar (CELMS) |
| A21 | *Pseudomonas resinovorans* |  | Elerin Toomik (CELMS) |
| L19 | *Rhodococcus erythropolis* |  | Elerin Toomik (CELMS) |
| A26 | *Rhodococcus erythropolis* |  | Elerin Toomik (CELMS) |
| A15 | *Rhodococcus erythropolis* |  | Elerin Toomik (CELMS) |
| A8 | *Rhodococcus erythropolis* |  | Elerin Toomik (CELMS) |
| A11B | *Cellulomonas* sp. |  | Elerin Toomik (CELMS) |
| A4X | *Bacillus subtilis* |  | Elerin Toomik (CELMS) |
| A24 | *Bacillus subtilis* |  | Elerin Toomik (CELMS) |
| L12ABA | *Bacillus simplex* |  | Elerin Toomik (CELMS) |
| A7 | *Bacillus pumilus* |  | Elerin Toomik (CELMS) |
| RP2 | *Rhodococcus* sp./*erythropolis* |  | Merike Jõessar (CELMS) |
| 43SK1 | *Kocuria rosea* |  | Eeva Heinaru, Sulev Kuuse, Julius Tarand (CELMS) |
| PC20 | *Pseudomonas fluorescens* | AY538264 (16S rRNA gene), DQ178200 (carA gene), AY887963 (Plasmid pNAH20), KY503036 (Plasmid pPHE20), KX893538 (Plasmid pG20) | Heinaru *et al.*, 2000 [15]; Eeva Heinaru (CELMS) |
| 6S4 | *Acinetobacter johnsonii* |  | Eeva Heinaru & Sulev Kuuse (CELMS) |
| P101 | *Pseudomonas thivervalensis* | MH061178 (Plasmid pPHE101), MN560131 (16S rRNA gene) | Elken *et al.*, 2020 [16]; Eeva Heinaru (CELMS) |
| KM2R4 | *Pseudomonas syringae* |  | Eeva Heinaru (CELMS) |
| VMR7 | *Pseudomonas stutzeri* |  | Eeva Heinaru (CELMS) |
| Phe16 | *Arthrobacter sulfonivorans* |  | Eeva Heinaru (CELMS) |

### Growth conditions and media

Cultures were incubated at 30 °C, and liquid cultures were shaken at 230 rpm. Rich, complex, undefined medium (Luria-Bertani medium, LB) was prepared using 10 g·L^-1^ tryptone, 10 g·L^-1^ NaCl, and 5 g·L^-1^ yeast extract. Wx/M9 minimal medium was based on Ariaki *et al*. (2020) [17] containing 5.7156 g·L^-1^ KH_2_PO_4_, 3.407 g·L^-1^ Na_2_HPO_4_, 1.0163 g·L^-1^ NH_4_Cl, 0.526 g·L^-1^ NaCl, and 1x trace elements solution containing 26.875 mg·L^-1^ MgO, 0.128 ml·L^-1^ HCl, 5 mg·L^-1^ CaCO_3_, 1.125 mg·L^-1^ FeSO_4_·7H_2_O, 3.6 mg·L^-1^ ZnSO_4_·7H_2_O, 2.8 mg·L^-1^ MnSO_4_·H_2_O, 0.625 mg·L^-1^ CuSO_4_·5H_2_O, 0.7 mg·L^-1^ CoSO_4_·7H_2_O, and 0.15 mg·L^-1^ H_3_BO_3_. Modified defined minimal medium (MMPAM) formulation was based on Borrero-de Acuña *et al*. (2021) [18] containing 6 g·L^-1^ KH_2_PO_4_, 7.09 g·L^-1^ Na_2_HPO_4_, 5 g·L^-1^ (NH_4_)_2_SO_4_, and 0.5 g·L^-1^ NaCl. The trace element composition remained the same.

### Plate reader growth assays

Precultures of *P. putida* KT2440 and PaW85 were made by inoculating a 5 mL LB medium with a single colony from an LB agar plate. The tubes were incubated and shaken at 30 °C and 230 rpm overnight in an orbital shaker. Half the culture volume was discarded and replaced with fresh LB medium. The cultures were then incubated and shaken for 90 minutes (approximately two doubling times) under the same parameters to ensure the cultures were in logarithmic growth phase. Osmotic stress tolerance assays were carried out using Wx/M9 medium supplemented with glucose or xylose at 2%, 4%, 6%, 8%, 10%, 12%, 15%, and 20% in 96-well clear polystyrene round-bottomed microplates with optically clear lids. pH tolerance assays were also performed using Wx/M9 medium supplemented with glucose at 6% concentration. The pH of the samples was adjusted to 7.1, 7.5, 7.7, 7.9, 8.1, 8.3, 8.7, 9.1, 9.5, and 10.0 with NaOH. The biomass of the precultures was measured for each strain to ensure the starting inoculates were the same and adjusted with sterile MilliQ water. 2 µL of inoculum was combined with 198 µL medium in each well. All microtiter assays were incubated at 30 °C in a BioTek Synergy MX plate reader, and A_600_ readings were taken every 15 minutes for 25 hours. Growth rates were calculated from triplicate samples from each plate reader experiment.

### Batch cultivations

Precultures were prepared under the same parameters as the plate reader growth assay except that Wx/M9 minimal medium at 2% glucose concentration was used. Precultures were inoculated into media with a starting A_600_ of 0.1. The medium-to-flask volume ratio was 1:5 in all experiments. The cultures were incubated and shaken at 30 °C and 230 rpm. Media composition varied depending on the experiment being run, and the cultures were periodically sampled to measure biomass (A_600_ was measured using a Hitachi U-1800 spectrophotometer), pH, total sugars, organic acids, and selected hydrolysis lignin (HL) monomer composition by HPLC. All cultures were grown in duplicate or triplicate.

### Simulated fed-batch cultivations

The simulated fed-batch media were prepared with components from EnPresso GmbH (Berlin, Germany). The components consisted of a powder-based polysaccharide added to the medium and an enzyme to release glucose from the polysaccharide over time. The enzymatic hydrolysis of the polysaccharide allowed for simulated carbon-limited cultivation. Precultures were prepared as described for the plate reader assays. A 200 g·L^-1^ stock EnPresso Pump 200 substrate solution was made using Wx/M9 medium and sterilized through a bottlecap filter (pore size 0.2 µm). Fractionated HL medium was supplemented with the stock solution to 20 g·L^-1^ in sterile flasks. The volume varied between 50 mL and 100 mL depending on the length of the cultivation and the number of samples to be collected. Enzyme reagent 0.6% was added at hour 4 or 6 post-inoculation, depending on the initial glucose concentration in the media and the doubling time under the culture environment. The A_600_ of the precultures was measured, and the flasks were inoculated to a working A_600_ of 0.1. Samples were taken periodically to measure the biomass, pH, and HL monomer composition by HPLC.

### Lignin fractionation

The reactive extrusion crude HL used in this study was obtained from Fibenol OÜ. The dried crude HL powder was used for further fractionation upon arrival from the biorefinery. HL fractionations were performed using Wx/M9 media alkalized with NaOH, KOH, or non-alkalized Wx/M9 medium. Alkalized Wx/M9 was adjusted to pH 11 with 10 M NaOH or KOH. The media were slowly combined with dry crude HL in a 6:1 ratio (w/w) and vigorously agitated for 45 minutes with a magnetic stir bar. The resultant slurry used in the batch cultivation was centrifuged at 110K rpm for 5 minutes, and the supernatant was filtered through a 0.2 µm bottlecap filter. The resultant slurries were autoclaved at 121 °C for 20 minutes and unfiltered for simulated fed-batch cultivations. EnPresso Pump 200 substrate solution was added at a final concentration of 20 g·L^-1^. Samples were taken before and after autoclaving and periodically during cultivation for analytics.

HL fractionation was also performed using the modified minimal media formulation MMPAM, previously described in the methods. The fractionation was performed in one 1 L bioreactor and agitated at 1000 rpm for 3 hours at 25 °C. The resultant slurry was centrifuged for 30 minutes at 4200 rpm, and the supernatant was filtered through a 0.2 µm bottlecap filter. The pH was subsequently normalized to 7.4 with 10 M NH_4_OH, and a 50 mL sample was taken for GC-MS analysis. The remaining volume was autoclaved at 121 °C for 20 minutes and used for analytics, simulated fed-batch cultivation, and strain screening experiments. This batch of fractionated HL medium is recorded as PAM511-0001.

HL was also fractionated for ion chromatography and spectrophotometry analyses using MilliQ H_2_O and Wx/M9. The ensuing slurries were autoclaved at 121 °C for 20 minutes, centrifuged at 110K rpm for 5 minutes, and the supernatant was filtered through a 0.2 µm bottlecap filter.

### Strain screening

Twenty-six bacterial strains were picked using a 1 µL sterile loop from R2A agar plates and cultivated at 30 °C and 150 rpm in 5 mL of the fractionated HL medium PAM511-0001. The growth medium was not supplemented with glucose. Samples were collected at hours 24 and 120 and centrifuged for 1 minute at 13K rpm. Two uninoculated media blanks (24h and 120h) were controls. The supernatants were stored at -20 °C and subsequently thawed for HPLC analysis to determine the HL monomer composition.

### HPLC analysis of samples

The monosaccharide and organic acid analyses were performed with the Shimadzu LC-2030C 3D Plus system using the method adapted from Monteiro de Oliveira *et al*. (2021) [19].

For monosaccharide detection, 20 µL of the sample was injected into a Rezex™ RPM-Monosaccharide Pb+2 (8%), LC column (300 × 7.8 mm) at 85 °C using a mobile phase of MilliQ H_2_O at a flow rate of 0.6 mL·min^-1^ and a Shimadzu RID-20A refractive index detector.

Organic acid detection analyses were performed by injecting 20 µL of sample into a Rezex™ ROA-Organic Acid H+ (8%) LC column (300 × 7.8 mm) at 45 °C using a mobile phase of 5 mM H_2_SO_4_ at a flow rate of 0.6 mL·min^-1^.

HL monomers were analyzed with the method adapted from Reyes-Rivera *et al*., (2015) [20]. 8 µL of sample was applied to a Kinetex® 2.6 µm C18 100 Å, LC column (150 × 4.6 mm) at 45 °C. The mobile phase of MilliQ H_2_O, methanol, and formic acid (80:20:0.16 v/v/v) flow rate was set to 0.6 mL·min^-1^, and monomers were detected with the UV-Vis detector at the wavelength of 265 nm.

### Ion chromatography and spectrophotometry

Analysis of the concentrations of chloride, nitrite, nitrate, phosphate, and sulfate was performed by injecting 20 µL of sample onto a Metrohm 930 Compact IC Flex system using a mobile phase of 0.1 M Na_2_CO_3_ and 0.1 M NaHCO_3_. Samples used to measure total nitrogen were prepared using the LATON® LCK138 kit, incubated in a Hach Lange LT200 instrument, and measured using a Hach Lange DR 2800 spectrophotometer (**Tab. S1**).

### Recording of UV-Vis spectra

A UV-Vis spectrophotometer (Thermo Scientific Evolution 160 using VisionLite 4.0 software) was used to measure the absorbance of the samples in the 200-600 nm wavelength range. Samples obtained from the various HL fractionations and the fermentations with *P. putida* strains were filtered and stored at -20 °C to prevent degradation or contamination. To measure the absorption spectra, appropriate dilutions were prepared using MilliQ water. Each sample was measured in triplicate quartz cuvettes with a 10 mm path length.

### Monomer and multimer analysis

The GC-MS analysis was performed similarly to Zhao *et al*., (2020) [21]. A 10 mL sample of fractionated crude HL was taken at hour 0 and hour 120 during a simulated fed-batch fermentation with *P. putida* KT2240. The sample was extracted with dichloromethane (4×10 mL), washed with saturated NH_4_Cl, dried with anhydrous MgSO_4_, and concentrated. The resulting oil products were acetylated using pyridine (1 mL) and acetic anhydride (1 mL) for 4 hours. The acetylated products were dissolved in 1 mL of dichloromethane and analyzed using the Agilent 7890A GC System with Agilent 5975 Inert XL MSD with Triple-Axis Detector and Agilent DB-5MS 30m × 0.250mm 0.25-micron column. GC-MS parameters included an inlet temperature of 300 °C, a total flow rate of Helium carrier gas at 21.1 mL·min^-1^, an ion source temperature of 250 °C, an interface temperature of 300 ºC, and an ion range of 45-800 (m/z). The oven program consisted of an initial temperature of 80 °C for 5 minutes, a ramp rate of 8 °C·min^-1^ up to 250 °C, a ramp rate of 5 °C·min^-1^ up to 300 °C, and a 30-minute hold at the final temperature, resulting in a total analysis time of 66.25 minutes.

## Results

### Physiological assessments

Characterizing the initial bacterial strains was essential. The objective was to ascertain the optimal growing conditions and to investigate how the strains reacted to different osmotic stress conditions. A physiological assessment was undertaken to achieve this, involving osmolarity experiments to determine the glucose and xylose tolerances of *P. putida* strains KT2440 and PaW85.

The glucose osmolarity microplate results revealed similar maximum specific growth rates (µ_max_) between 20 and 60 g·L^-1^, and neither strain could tolerate 200 g·L^-1^ glucose concentration (**Fig. 1a**).

**Figure 1.**
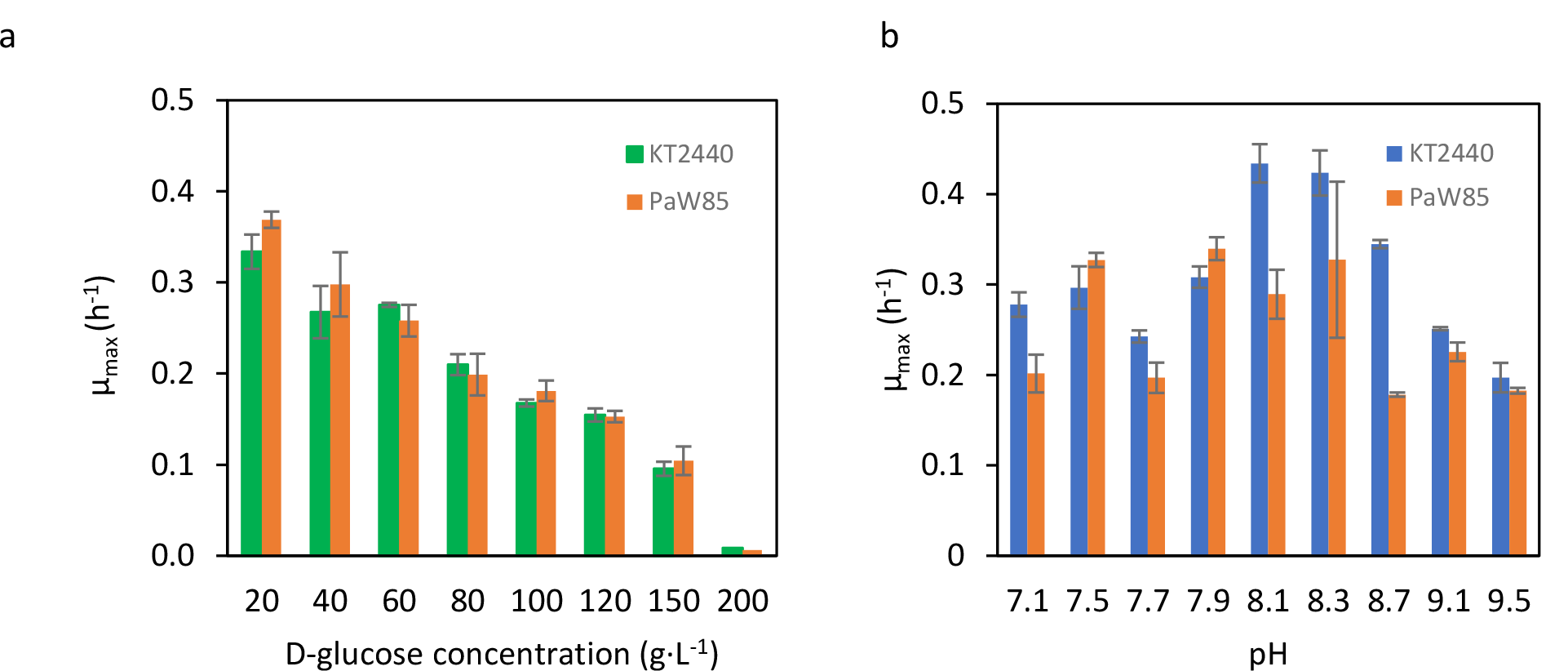
Glucose osmolarity and pH tolerance profiles for *P. putida* KT2440 and PaW85. Maximum specific growth rates at various glucose concentrations in Wx/M9 minimal medium at 30 °C pH 7.2 over 25 hours (**a**). Maximum specific growth rates at various pH levels in Wx/M9 minimal medium at 30 °C and 60 g·L^-1^ glucose, pH adjusted with NaOH (**b**). Standard error was used for error bars.

The xylose osmolarity experiments revealed that strains KT2440 and PaW85 could not grow in a medium containing xylose as the only carbon source (**Fig. S2**).

Another physiological evaluation conducted to determine the ideal growth conditions for the KT2440 and PaW85 strains involved a pH tolerance investigation. The results revealed that both strains exhibited a significantly broader pH tolerance range than initially expected. Notably, robust growth was observed at elevated pH levels, up to 9.5 (**Fig. 1b**).

### Growth in fractionated HL

HL fractionation was initially performed using NaOH-alkalized Wx/M9 minimal medium. The resulting solution underwent sterile filtration and was used for initial batch cultivation with *P. putida* KT2440. Samples were collected at regular 24-hour intervals and subjected to HPLC analysis to determine the sugar concentration. The outcomes of this cultivation indicated complete consumption of glucose and xylose within the initial 24 hours, with a peak A_600_ of approximately 5 (**Fig. 2**). Moreover, the medium exhibited an absorbance peak at 280 nm prior to cultivation, which decreased by hour 48 indicating a change in the dissolved aromatic compounds (**Fig. 3**).

**Figure 2.**
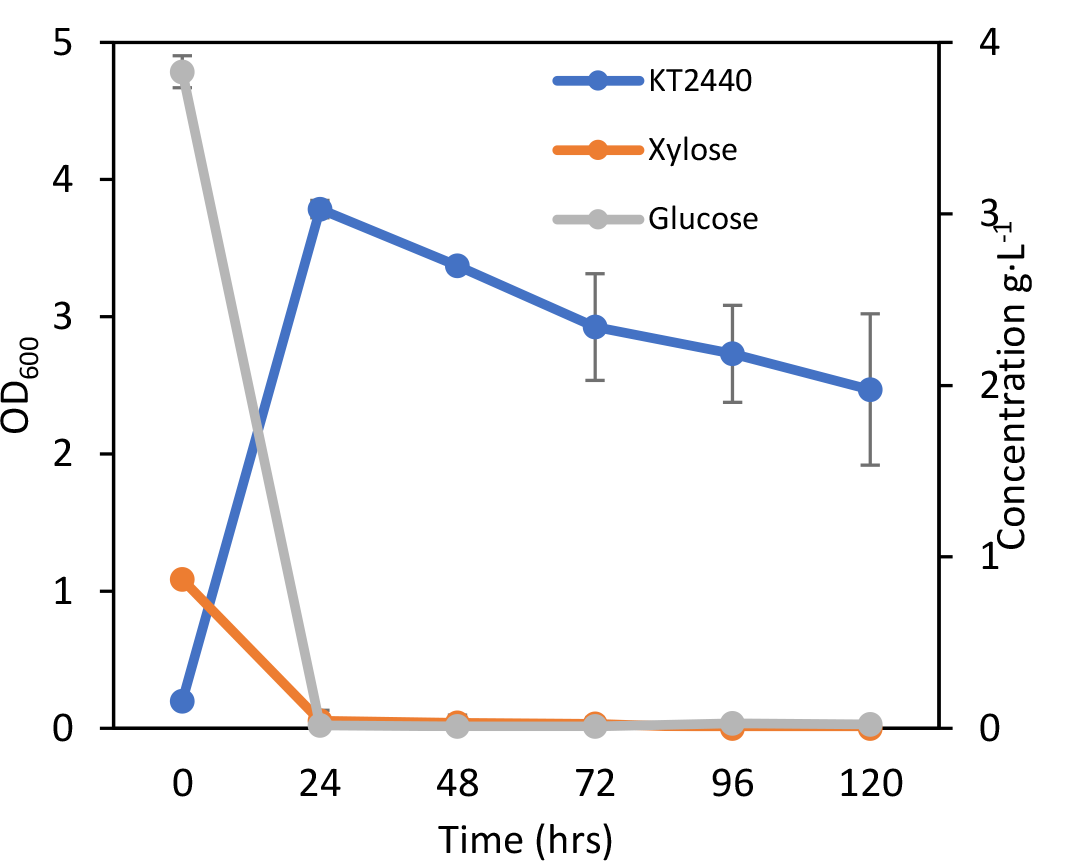
Glucose and xylose consumption of *P. putida* KT2440. Growth curve and glucose and xylose utilization during 120-hour batch fermentation in filtered Wx/M9-fractionated HL medium. Standard error was used for error bars.

**Figure 3.**
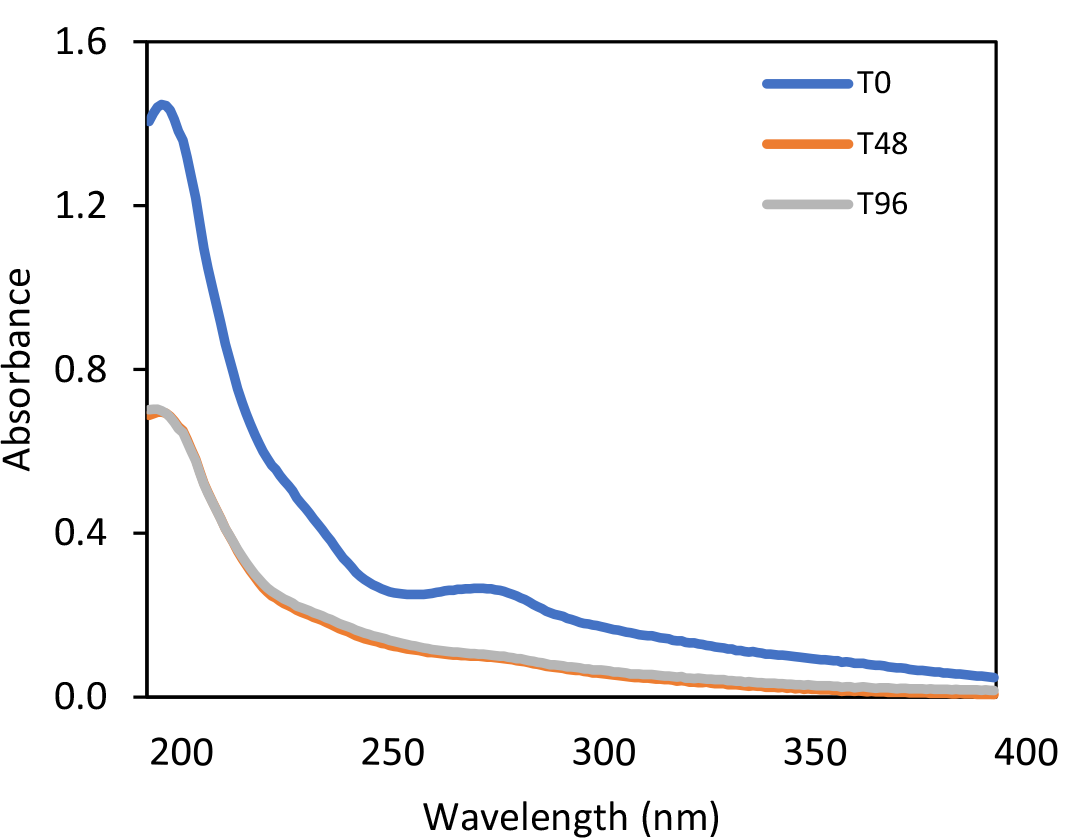
Absorbance spectra of fractionated HL medium. Absorbance spectra of filtered NaOH-alkalized HL medium at hours 0, 48, and 96 of cultivation with *P. putida* KT2440. The wavelength range is 200-400 nm and dilution 400X.

### Fractionated HL simulated fed-batch cultivation

The simulated fed-batch cultivation technique involves maintaining carbon-starved conditions for the bacterial culture throughout the fermentation process. The HL was fractionated using Wx/M9 minimal medium alkalized with NaOH and KOH. Glucose and xylose were initially measured to be approximately 4 g·L^-1^ and 2 g·L^-1^, respectively. A polysaccharide was added at a concentration of 20 g·L^-1^, and an enzyme reagent was added at 0.6% concentration four hours after the start of the fermentation. Thus, KT2440 had a steady supply of glucose during the fermentation due to the enzymatic breakdown of the polysaccharide into glucose.

Changes in the HL monomer concentrations were analyzed via HPLC. The results showed that over 0.5 g·L^-1^ of catechol was solubilized from the fractionation. KT2440 completely catabolized the catechol within the first 4 hours, which remained nearly undetectable throughout the experiment. In contrast, the CCMA concentration increased approximately sevenfold within the first 24 hours and then stabilized for the rest of the fermentation (**Fig. 4a**). Over time, the samples lightened in color (**Fig. 4b**), and the odor changed from nutty to slightly acidic by hour 96.

**Figure 4.**
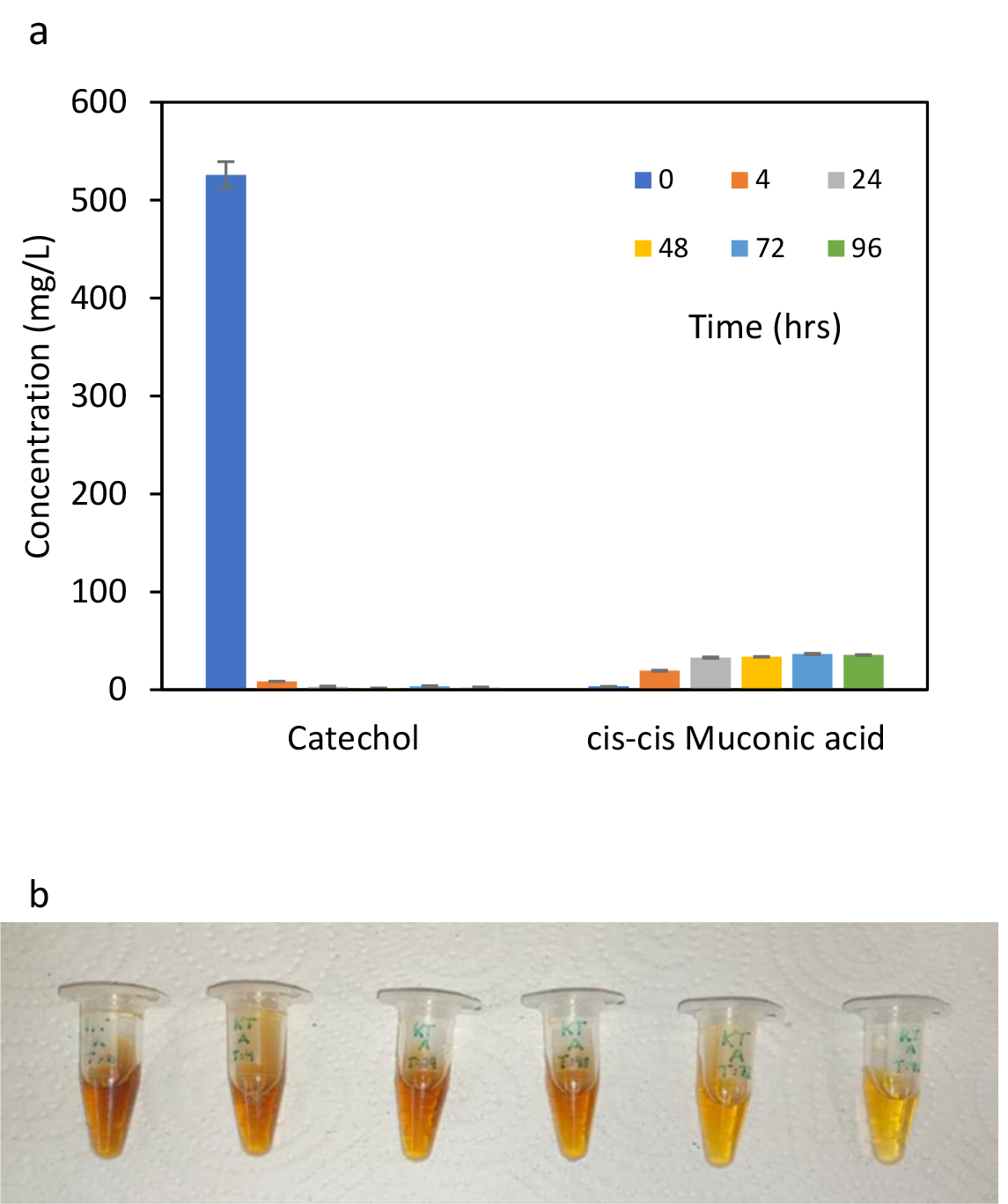
Catechol and CCMA concentrations and sample color changes in filtered HL cultivation. Variation of catechol and *cis,cis*-muconic acid concentrations in filtered Wx/M9-fractionated HL medium during 96-hour simulated fed-batch fermentation with *P. putida* KT2440 (**a**). Standard error was used for error bars. Snapshot of samples taken during 96-hour simulated fed-batch fermentation of P. putida KT2440 in Wx/M9-fractionated HL medium (**b**).

### HL slurry cultivations

This simulated fed-batch cultivation used an HL slurry as the growth medium for *P. putida* KT2440. Dry HL was fractionated using two alkaline Wx/M9 media solutions and one non-alkaline medium. The solids were not removed. Between 24 and 48 hours, the glucose concentration decreased significantly, coinciding with a drop in pH. The glucose concentration subsequently increased, indicating ongoing hydrolysis of the polysaccharide (**Fig. S3a**).

Moreover, there is an increase in citric acid production until hour 48, after which levels stabilize, indicating the cultures did not tolerate acidic conditions below pH 5 (**Fig. S3b**). HL monomer concentrations of the samples were assessed using HPLC. In the NaOH and KOH-alkalized HL cultivations, catechol was nearly completely catabolized within 4 hours, leading to an initial increase in CCMA concentration, which stabilized after 48 hours. However, the initial catechol concentration was significantly lower than expected in both cases (**Fig. S4**).

### Ion chromatographic analysis of fractionated HL

The ion concentrations in water-fractionated and Wx/M9-fractionated HL were compared to determine the extent of ion dissolution during fractionation and the necessity of fractionation with a defined minimal medium. The findings revealed that approximately 75% of dissolved phosphate originated from the HL in the Wx/M9 fractionation. Nitrate and nitrite levels were similar in both samples, while the Wx/M9 fractionation exhibited higher total nitrogen due to dissolved NH_4_OH and components of the medium. Chloride and sulfate levels were also significantly elevated in the Wx/M9 fractionation (**Tab. S1**).

### Multimeric to monomeric shift in alkaline HL fermentations

Zhao *et al*., (2020) [21] conducted a study wherein common lignin-derived trimers, dimers, and monomers were synthesized and analyzed using GC-MS. This study used the same Selected Ion Monitoring (SIM) mode parameters to monitor specific mass-to-charge ratios of acetylated target lignin compounds present in the liquid phase when fractionating HL under alkaline conditions. Samples were obtained at hours 0 and 120 of a simulated fed-batch cultivation with *P. putida* KT2440 and followed by extraction and acetylation according to the protocol in publication by Zhao. The resulting chromatograms show a shift to shorter acquisition times, which correspond to lignin monomers, compared to the latter sample with no peaks within the 32-to-44-minute range (**Fig. 5a, b**). These results indicate that *P. putida* KT2440 can potentially consume lignin oligomers and convert them into monomeric lignin-derived compounds.

**Figure 5.**
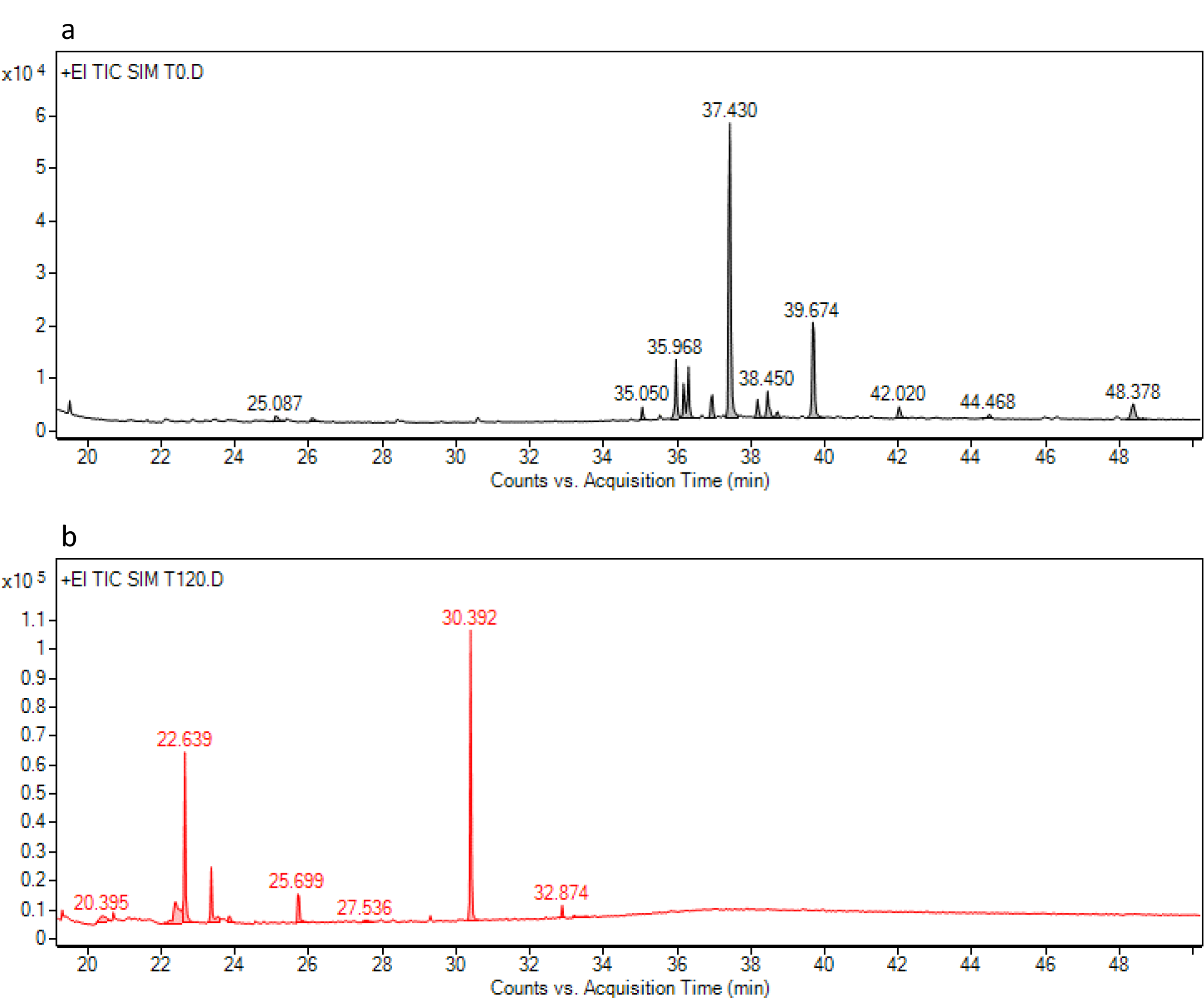
Acquisition time of HL-derived compounds. Chromatogram of acetylated HL-derived di/trimers at hour 0 of alkalized HL fractionation sample from simulated fed-batch cultivation with *P. putida* KT2440 (**a**). Chromatogram of acetylated HL-derived monomers at hour 120 of alkalized HL fractionation sample from simulated fed-batch cultivation with *P. putida* KT2440 (**b**).

### Strain screening

The HPLC data from batch cultivations of 26 strains isolated from soil and water samples was analyzed at different time points to identify potential candidates for future HL research. Five compounds were identified that were above the stochastic range.

By hour 24, most strains catabolized over 75% of the catechol. CCMA increased over 50% for *P. putida* KT2440, *P. putida* 1S7, *Pseudomonas* sp./*defluvii* VAS4, *P. resinovorans* A21, and *P. putida* PJK7. 5-hydroxymethylfurfural (5-HMF) concentrations decrease significantly at hour 24 and hour 120 for most strains. An exception is *B. simplex* L12ABA, in whose samples 5-HMF remains relatively unchanged. PCA concentrations increase the most in strains *Enterobacter* sp./*Lelliottia* sp. T13 and *M. chitosanitabida* 1R2A4 at hours 24 and 120. Regarding syringic acid, most screened strains show a slight decrease in concentration at 24 hours, except *P. putida* KT2440 and *M. chitosanitabida* 1R2A4, which show little change in concentration. Conversely, *P. putida* 1S7 and *Enterobacter* sp./*Lelliottia* sp. T13 catabolize the most syringic acid at hours 24 and 120. Regarding vanillic acid, the concentrations increased by over 25% in *Pseudomonas* sp./*defluvii* VAS4, *P. putida* PJK7, and *R. erythropolis* A26 in 24 hours. Contrarily, strains *P. putida* 1S7 and *Enterobacter* sp./*Lelliottia* sp. T13 catabolize nearly all vanillic acid within 24 hours (**Tab. 2**).

**Table 2.**
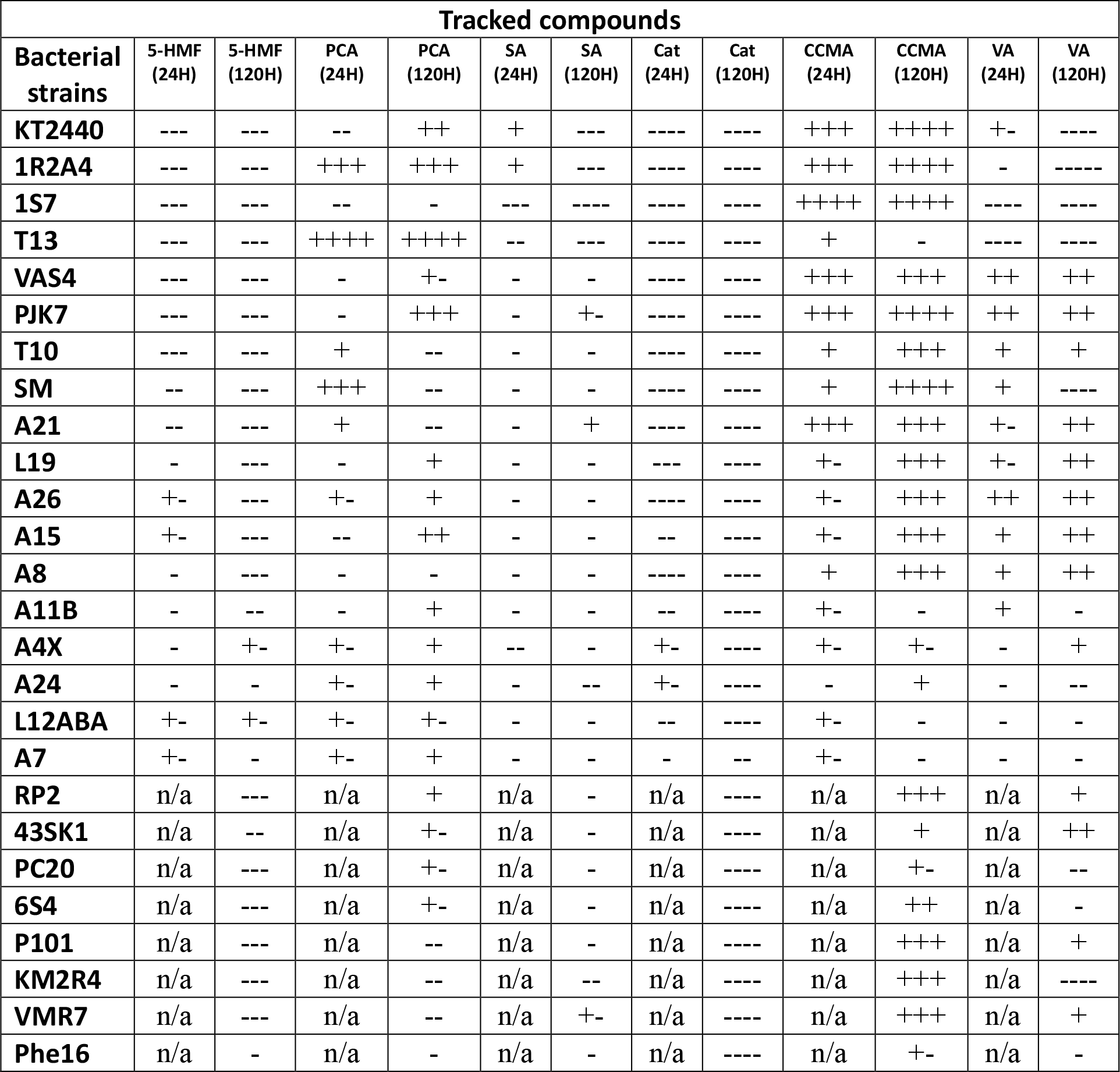
Bacterial strain screening. Conversion of lignin-derived compounds after 24 and 120 hours of bacterial cultivation in MMPAM-fractionated HL medium compared to the contents in uninoculated 24-hour and 120-hour controls. The compounds are labeled as follows: 5-HMF: 5-hydroxymethylfurfural; PCA: protocatechuic acid; SA: syringic acid; Cat: catechol; CCMA: *cis,cis*-muconic acid; VA: vanillic acid. The symbols in the table indicate the percentage range of change in compound concentration. ----: >75% decrease; ---: 50-75% decrease; --: 25-50% decrease; -: 5-25% decrease; +-: 5% decrease to 5% increase; +: 5-35% increase; ++: 25-50% increase; +++:50-75% increase; ++++: >75% increase.

| Tracked compounds |  |  |  |  |  |  |  |  |  |  |  |  |
| --- | --- | --- | --- | --- | --- | --- | --- | --- | --- | --- | --- | --- |
| Bacterial strains | 5-HMF (24H) | 5-HMF (120H) | PCA (24H) | PCA (120H) | SA (24H) | SA (120H) | Cat (24H) | Cat (120H) | CCMA (24H) | CCMA (120H) | VA (24H) | VA (120H) |
| KT2440 | --- | --- | -- | ++ | + | --- | ---- | ---- | +++ | ++++ | +/- | ---- |
| 1R2A4 | --- | --- | +++ | +++ | + | --- | ---- | ---- | +++ | ++++ | - | ---- |
| 1S7 | --- | --- | -- | - | --- | ---- | ---- | ---- | ++++ | ++++ | ---- | ---- |
| T13 | --- | --- | ++++ | ++++ | -- | --- | ---- | ---- | + | - | ---- | ---- |
| VAS4 | --- | --- | - | +/- | - | - | ---- | ---- | +++ | +++ | ++ | ++ |
| PJK7 | --- | --- | - | +++ | - | +/- | ---- | ---- | +++ | ++++ | ++ | ++ |
| T10 | --- | --- | + | -- | - | - | ---- | ---- | + | +++ | + | + |
| SM | -- | --- | +++ | -- | - | - | ---- | ---- | + | ++++ | + | ---- |
| A21 | -- | --- | + | -- | - | + | ---- | ---- | +++ | +++ | +/- | ++ |
| L19 | - | --- | - | + | - | - | --- | ---- | +/- | +++ | +/- | ++ |
| A26 | +/- | --- | +/- | + | - | - | ---- | ---- | +/- | +++ | ++ | ++ |
| A15 | +/- | --- | -- | ++ | - | - | -- | ---- | +/- | +++ | + | ++ |
| A8 | - | --- | - | - | - | - | ---- | ---- | + | +++ | + | ++ |
| A11B | - | -- | - | + | - | - | -- | ---- | +/- | - | + | - |
| A4X | - | +/- | +/- | + | -- | - | +/- | ---- | +/- | +/- | - | + |
| A24 | - | - | +/- | + | - | -- | +/- | ---- | - | + | - | -- |
| L12ABA | +/- | +/- | +/- | +/- | - | - | -- | ---- | +/- | - | - | - |
| A7 | +/- | - | +/- | + | - | - | - | -- | +/- | - | - | - |
| RP2 | n/a | --- | n/a | + | n/a | - | n/a | ---- | n/a | +++ | n/a | + |
| 43SK1 | n/a | -- | n/a | +/- | n/a | - | n/a | ---- | n/a | + | n/a | ++ |
| PC20 | n/a | --- | n/a | +/- | n/a | - | n/a | ---- | n/a | +/- | n/a | -- |
| 6S4 | n/a | --- | n/a | +/- | n/a | - | n/a | ---- | n/a | ++ | n/a | - |
| P101 | n/a | --- | n/a | -- | n/a | - | n/a | ---- | n/a | +++ | n/a | + |
| KM2R4 | n/a | --- | n/a | -- | n/a | -- | n/a | ---- | n/a | +++ | n/a | ---- |
| VMR7 | n/a | --- | n/a | -- | n/a | +/- | n/a | ---- | n/a | +++ | n/a | + |
| Phe16 | n/a | - | n/a | - | n/a | - | n/a | ---- | n/a | +/- | n/a | - |

## Discussion

*P. putida* KT2440 exhibited broad pH tolerance and robust growth at elevated pH levels. Its high pH tolerance is promising because HL is only water soluble in alkaline conditions [22], which was validated by UV-spectroscopy analysis. Given that *P. putida* can thrive in alkaline conditions, using alkali-fractionated HL – containing abundant dissolved HL – as an effective growth medium proved possible. Furthermore, the complete consumption of sugars within 24 hours during the initial batch fermentation and the marked decrease in the absorbance peak at 280 nm demonstrated, for the first time, the viability of fractionated HL as a substrate for *P. putida* cultivation for the production of value-added chemicals such as *cis,cis*-muconic acid. However, the decrease in xylose concentration was unexpected. Xylose assimilation occurs via an isomerase or two oxidative metabolic pathways in microbes, such as *P. taiwanensis* VLB120 [23], but *P. putida* KT2440 cannot metabolize xylose [24].

HL is an adequate phosphate source for this study’s simulated fed-batch growth conditions but lacks nitrogen and other essential trace ions. Consequently, water fractionation is not feasible for supporting bacterial growth without an additional nitrogen source and trace metals.

Results from the simulated fed-batch fermentation indicate that when catechol concentration is high, CCMA will be produced more quickly than degraded, which is a promising prospect for fed-batch or continuous cultivation experiments while feeding HL media preparations.

Filtered alkaline HL fractionations contained over double the amount of dissolved catechol as unfiltered ones. The reasons for this are not entirely understood; however, catechol is unstable and is easily oxidized into highly reactive quinones in an aerated aqueous solution at neutral to alkaline pH [25,26]. Although the filtered alkaline fractionation is likely the better method for CCMA production, the unfiltered fractionation may be better for producing other compounds and would be easier to upscale.

GC-MS results indicate a possible conversion of HL-derived multimers into monomers during the fermentation process with KT2440, suggesting *P. putida* may have produced extracellular enzymes, depolymerizing some of the detected dimers and trimers in solution.

Five of the screened bacterial strains exhibited rapid catechol catabolism and the highest production of CCMA, making them promising candidates for enzyme discovery and genetic engineering: *P. putida* KT2440, *P. putida* 1S7, *P. putida* PJK7, *Pseudomonas* sp.*/defluvii* VAS4, and *Pseudomonas resinovorans* A21. Of these, all except *P. resinovorans* A21 showed glucose and xylose metabolism. The discovery of efficient enzymes in new strains could eventually enhance CCMA production in *P. putida*, lowering the marginal costs of renewable plastics production. Moreover, *Rhodococcus erythropolis* A26, *P. putida* PJK7, and *Pseudomonas* sp.*/defluvii* VAS4 showed the most potential for vanillic acid production, which remains in high demand in the pharmaceutical, cosmetic, and food industries. However, it is notable that the screening process may have a cultivation bias towards *Pseudomonas* strains due to the modified MMPAM minimal medium, which was optimized for *P. putida* when creating the PAM511-0001 fractionated HL batch on which the strains were screened.

## Conclusions

We demonstrated that hydrolysis lignin is a viable substrate for *Pseudomonas putida* cultivation to produce value-added chemicals and that alkaline fractionation methods are practicable due to the strains’ high pH tolerances. Filtered NaOH and KOH fractionations dissolve more catechol and are more effective for CCMA production than similar unfiltered fractionations, and *P. putida* KT2440 may be capable of enzymatically depolymerizing HL-derived multimers in solution. Several screened strains show potential for enzyme discovery, allowing for the modification or replacement of enzymatic pathways in *P. putida* with more efficient enzymes from current and future screened strains.

## List of abbreviations

(HL): Hydrolysis lignin
(CCMA): *cis,cis*-muconic acid
(GHG): greenhouse gas
(BDP): benzoate degradation pathway
(LB): Luria broth
(PCA): protocatechuic acid
(MMPAM): modified minimal media formulation
(SIM): Selected Ion Monitoring
(GC-MS): gas chromatography-mass spectrometry
(HPLC): high-performance liquid chromatography

## Declarations

**Ethics approval and consent to participate**

**Consent for publication**

**Availability of data and material**

**Competing interests**

## Funding

ERDF and the Estonian Research Council supported this study via projects RESTA6 and RESTA22

## Authors’ contributions

PM, HV, CM, SV, MJ, SB, SS performed experiments. SV, MJ, SB, SS, MK, ML designed experiments and edited the manuscript. PM, HV, SB, SS performed GC-MS analysis. PM, CM, SB performed LC and IC analysis. PM, SV, MJ, SB performed the strain screening. KV and ML conceived the project. SS, KV, and ML funded the project. All authors read, edited, and approved the manuscript.

## Authors’ information

## Supplementary materials

**Table S1.**
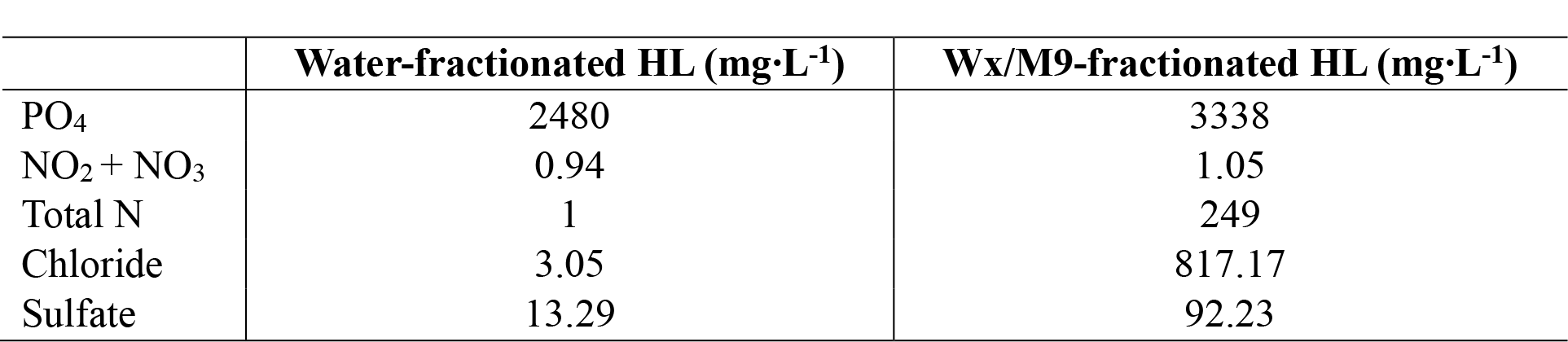
Phosphate, nitrogen, chloride, and sulfate concentrations in fractionated HL.

**Figure S2.**
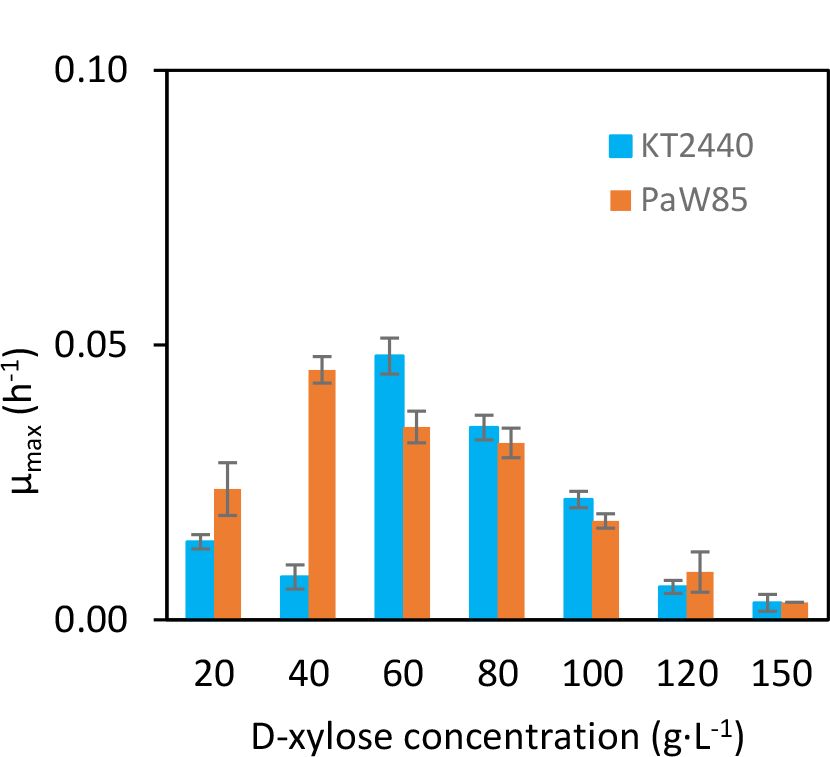
Xylose osmolarity profiles for *P. putida* KT2440 and PaW85. Maximum specific growth rates at various xylose concentrations in Wx/M9 minimal medium at 30 °C pH 7.2 over 25 hours. Standard error was used for error bars.

**Figure S3.**
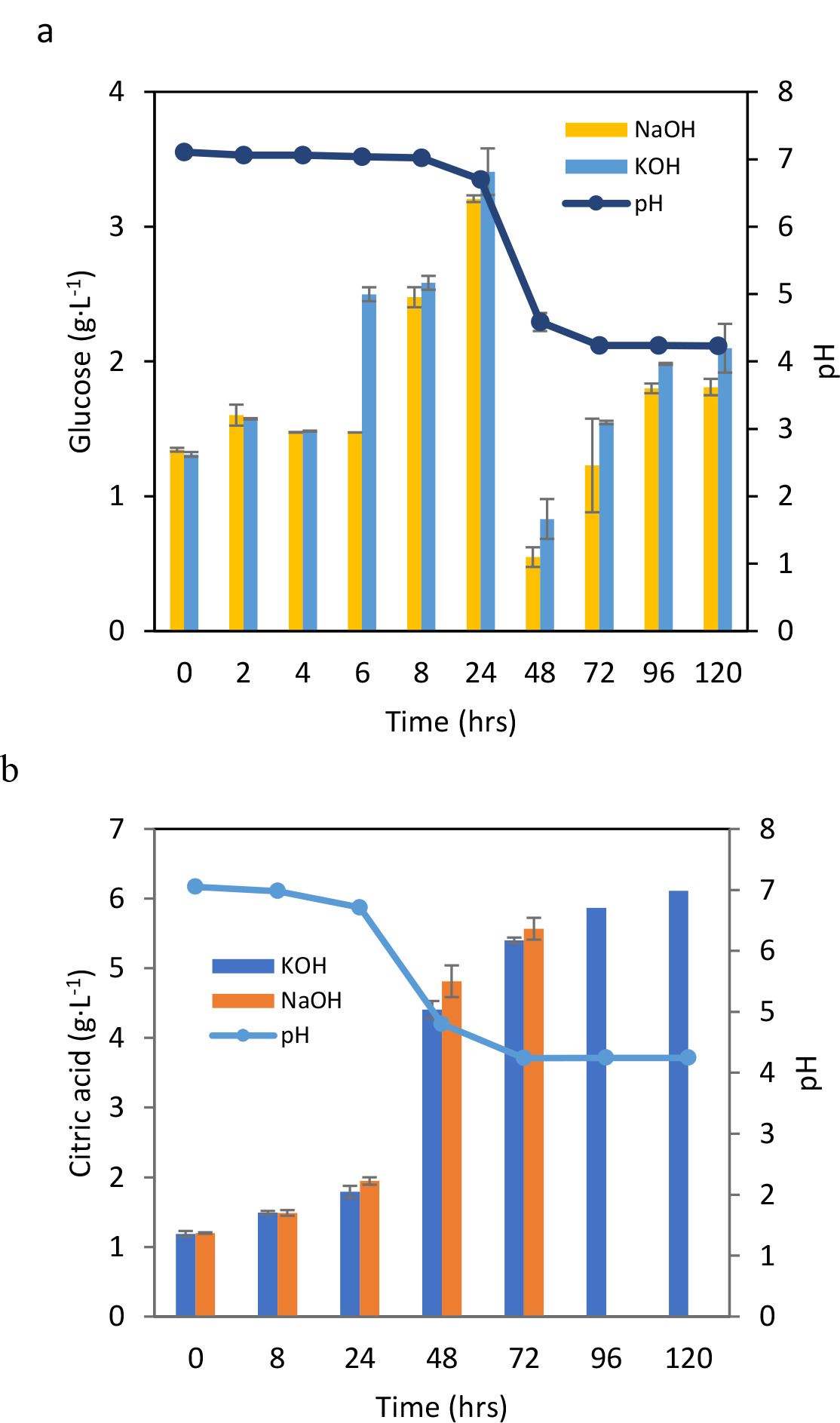
Glucose and citric acid concentration dynamics and pH level changes. 120-hour simulated fed-batch fermentation with P. putida KT2440 in unfiltered NaOH-and KOH-alkalized HL media. Glucose concentration dynamics (**a**). Citric acid concentration dynamics (**b**). Standard error was used for error bars.

**Figure S4.**
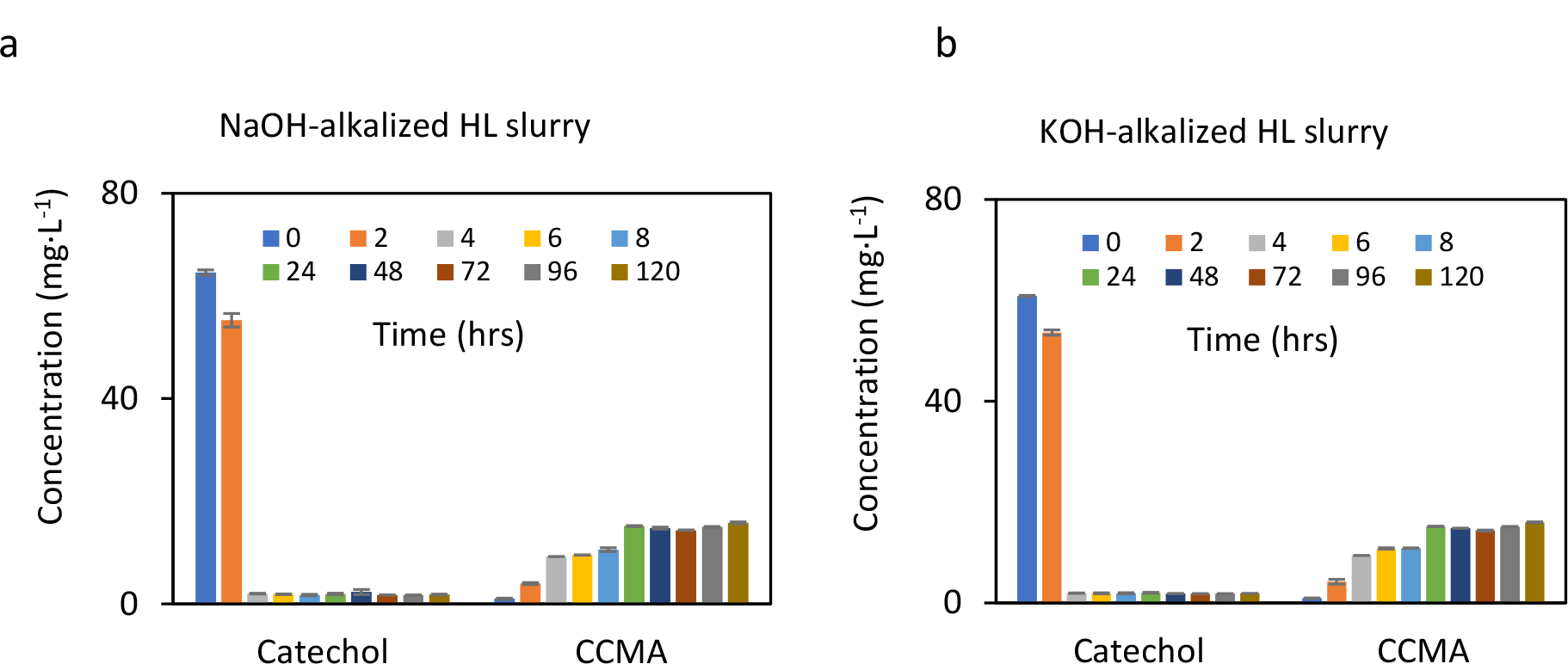
Changes in the concentrations of catechol and *cis,cis*-muconic acid. 120-hour simulated fed-batch fermentation of unfiltered HL fractionations made with Wx/M9 minimal medium and alkalized with NaOH (**a**) and KOH (**b**). Standard error was used for the error bars.

## Notes

### Competing Interest Statement

The authors have declared no competing interest.

