## Supplemental Information for "Conversion of aromatic compounds from fractionated industrial hydrolysis lignin by *Pseudomonas putida* and environmental microbial strains"

**Supplementary materials**

**Table S1. Phosphate, nitrogen, chloride, and sulfate concentrations in fractionated HL**

| | Water-fractionated HL (mg·L$^{-1}$) | Wx/M9-fractionated HL (mg·L$^{-1}$) |
|---|---|---|
| $PO_4$ | 2480 | 3338 |
| $NO_2 + NO_3$ | 0.94 | 1.05 |
| Total N | 1 | 249 |
| Chloride | 3.05 | 817.17 |
| Sulfate | 13.29 | 92.23 |

**Figure S2. Xylose osmolarity profiles for *P. putida* KT2440 and PaW85**

Maximum specific growth rates at various xylose concentrations in Wx/M9 minimal medium at 30 °C pH 7.2 over 25 hours. Standard error was used for error bars.

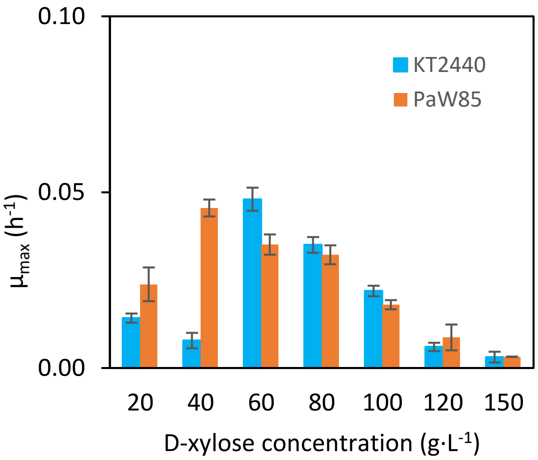

**Figure S3. Glucose and citric acid concentration dynamics and pH level changes**

120-hour simulated fed-batch fermentation with P. putida KT2440 in unfiltered NaOH-and KOH-alkalized HL media. Glucose concentration dynamics (**a**). Citric acid concentration dynamics (**b**). Standard error was used for error bars.

a

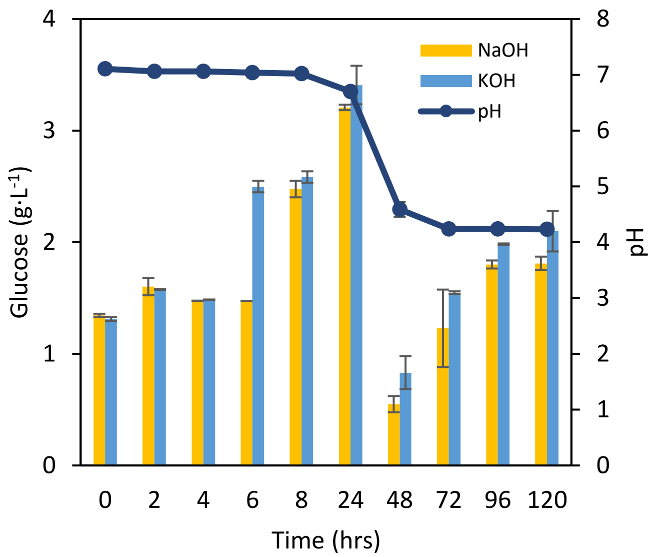


b

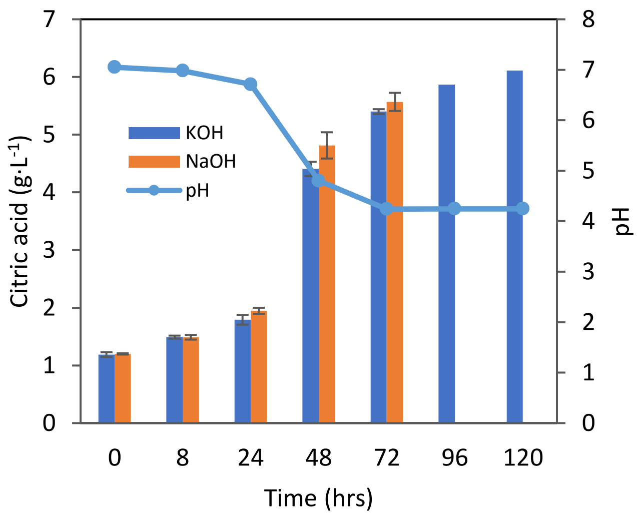

**Figure S4. Changes in the concentrations of catechol and *cis,cis*-muconic acid**

120-hour simulated fed-batch fermentation of unfiltered HL fractionations made with Wx/M9 minimal medium and alkalized with NaOH (**a**) and KOH (**b**). Standard error was used for the error bars.

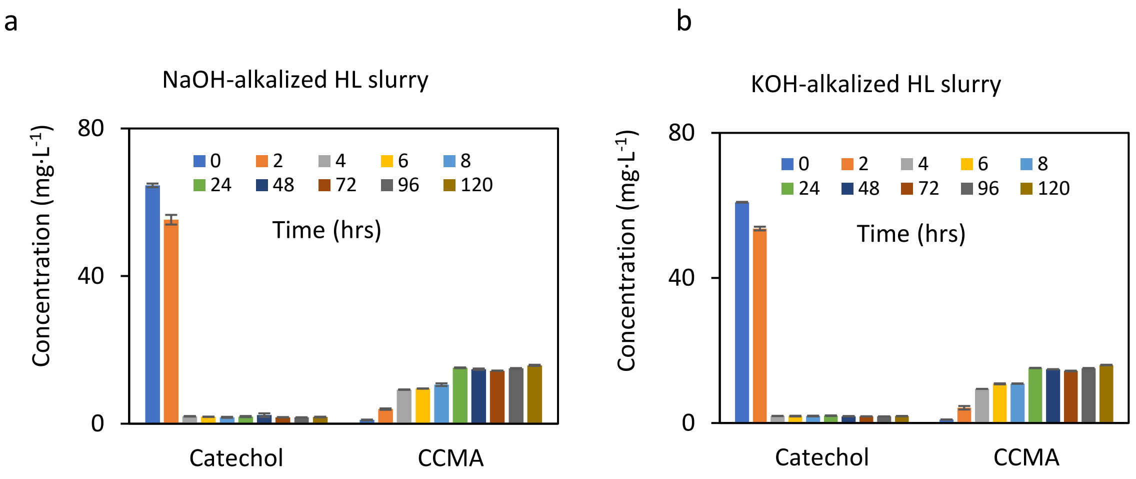
